# Microenvironment of metastasis reveals key predictors of PD-1 blockade response in renal cell carcinoma

**DOI:** 10.1101/2023.07.17.548676

**Authors:** Florian Jeanneret, Pauline Bazelle, Sarah Schoch, Catherine Pillet, In Hwa Um, Assilah Bouzit, Bertrand Evrard, Evan Seffar, Frédéric Chalmel, Javier A Alfaro, Catia Pesquita, Fabio Massimo Zanzotto, Mark Stares, Stefan N Symeonides, Alexander Laird, Jean-Alexandre Long, Jean Luc Descotes, Delphine Pflieger, David J Harrison, Odile Filhol, Håkan Axelson, Christophe Battail

**Affiliations:** Univ. Grenoble Alpes, IRIG, Laboratoire Biosciences et Bioingénierie pour la Santé, UA 13 INSERM-CEA-UGA, CNRS FR 2048; 38000 Grenoble, France; Division of Translational Cancer Research, Department of Laboratory Medicine Lund University; Lund, Sweden; Univ. Grenoble Alpes, IRIG, Biosanté, UMR 1292 INSERM-CEA-UGA; 38000 Grenoble, France; School of Medicine, University of St Andrews; North Haugh, St Andrews KY16 9TF, Scotland, UK; Centre hospitalier universitaire Grenoble Alpes, CS 10217; 38043 Grenoble cedex 9, France; Univ Rennes, Inserm, EHESP, Irset (Institut de recherche en santé, environnement et travail) - UMR_S 1085; F-35000 Rennes, France; Marie-Josée and Henry R. Kravis Center for Molecular Oncology, Memorial Sloan Kettering Cancer Center, New York, NY, USA. Department of Epidemiology and Biostatistics, Memorial Sloan Kettering Cancer Center; New York, NY, USA; International Centre for Cancer Vaccine Science, University of Gdansk; Gdansk, Poland; LASIGE, Faculdade de Ciências, Universidade de Lisboa; Lisboa, Portugal; Department of Enterprise Engineering, University of Rome “Tor Vergata”; Rome, Italy; Edinburgh Cancer Centre, Western General Hospital, NHS Lothian; Edinburgh, UK; Institute of Genetics and Cancer, Cancer Research UK Edinburgh Centre, University of Edinburgh; Edinburgh, UK; Department of Urology, Western General Hospital, NHS Lothian; Edinburgh, UK

## Abstract

Immune checkpoint blockade (ICB) therapies have improved the overall survival (OS) of many patients with advanced cancers. However, the response rate to ICB varies widely among patients, exposing non-responders to potentially severe immune-related adverse events. The discovery of new biomarkers to identify patients responding to ICB is now a critical need in the clinic. We therefore investigated the tumor microenvironment (TME) of advanced clear cell renal cell carcinoma (ccRCC) samples from primary and metastatic sites to identify molecular and cellular markers of response to ICB. We revealed a significant discrepancy in treatment response between subgroups based on cell fractions inferred from metastatic sites. One of the subgroups was enriched in non-responders and harbored a lower fraction of CD8+ T cells and plasma cells, as well as a decreased expression of immunoglobulin genes. In addition, we developed the Tumor-Immunity Differential (TID) score which combines features from tumor cells and the TME to accurately predict response to anti-PD-1 immunotherapy (AUC-ROC=0.88, log-rank tests for PFS P < 0.0001, OS P = 0.01). Finally, we also defined TID-related genes (*YWHAE*, *CXCR6* and *BTF3*), among which *YWHAE* was validated as a robust predictive marker of ICB response in independent cohorts of pre- or on-treatment biopsies of melanoma and lung cancers. Overall, these results provide a rationale to further explore variations in the cell composition of metastatic sites, and underlying gene signatures, to predict patient response to ICB treatments.

**One Sentence Summary:** Tumor microenvironment balance of metastasis and associated genes are key predictors of immunotherapy patient response in kidney cancer.

## INTRODUCTION

Immune checkpoint blockage (ICB) therapies have shown remarkable success in decreasing the recurrence rate and improving the overall survival of patients with metastatic clear cell Renal Cell Carcinoma (ccRCC) (*1–7*). However, only a subset of patients experiences long-term clinical response with these therapies. This diversity in response to ICB not only underscores the need for novel predictive markers to identify patients who will benefit from these treatments, but also the need to reveal molecular and cellular mechanisms involved in the acquisition of therapeutic resistance.

Tumor mutation burden (TMB), an established marker of response to immunotherapy in various cancers (*8*), was shown not to correlate with patient prognostics in ccRCC (*9–11*). Expression of immune checkpoint genes, such as PD-1 and PD-L1, in cancer cells and in tumor-infiltrating cells is another feature associated with favorable response to ICB in several cancer types (*12*). However, in ccRCC a low percentage of samples express PD-1 and PD-L1 (*13*) an association between their expression level and ccRCC aggressiveness and prognosis remains controversial (*5, 14–17*). For PD-L1, sources of variability in its expression were found to be related to intratumoral heterogeneity, differences between primary and metastatic sites and the lack of standardization of experimental techniques for measuring abundances (*18*).

Beyond single gene markers, predictive scores based on gene expression signatures have shown promising results in predicting ICB response for various cancer types. Among them, the 15 immune gene score IMPRES (*19*), the 18 IFN-γ-responsive gene score GEP (*20*), MHC-I/MHC-II (*21*) and MIAS (*22*) scores demonstrated a strong predictive power of response to ICB therapies in metastatic melanoma but were not evaluated in ccRCC. The JAVELIN score was specifically developed and tested on kidney cancer (*1*). Nevertheless, although they carry some predictive capacities, to our knowledge none of these scores have yet been used in clinical practice (*23, 24*).

The immune components and composition of the tumor microenvironment (TME) have been extensively described to be part of the major determinants of the response to ICB (*25–27*). For example, the level of CD8+ infiltrating T cells has been associated with a good prognosis in many cancer types (*25*) whereas the role of tumor infiltrating CD8+ T cells in survival of patients with ccRCC remains controversial. The response to ICB may also depend on the spatial localization of immune cells. For instance, tertiary lymphoid structures (TLS) composed of mature dendritic cells, B-cells and CD8+ T-cell have been reported to be important in predicting ICB response in ccRCC (*28–31*). Regarding B cells, their location in mature TLS has been suggested to mediate antibody release, T cells activation and has been described to be enriched in good ICB responders in ccRCC patients (*32*). Tumor-associated macrophages (TAM) were found to contribute to ccRCC progression and metastasis and to mediate immunosuppression Thus, the factors driving ICB resistance remain largely to be deciphered and effective markers to predict ICB response for ccRCC in clinical practice are seriously lacking.

In this work, we perform an extensive relabeling of single-cell RNA-seq data from ccRCC into more than 20 cell types and assess their robustness and accuracy in predicting TME composition by cell deconvolution of bulk RNA-seq data derived from patients in ICB clinical trials (anti-PD-1). The proportions of these reference cell types predicted from advanced ccRCC transcriptomes of primary and metastatic sites separately reveal new key cellular and molecular markers of ICB response. We highlight the biological value provided by the metastatic sites by identifying three TME profiles with significantly different clinical outcomes. In particular, the CD8+ T cells and plasma cells interplay is a major factor influencing the PD1 blockade response of patients with ccRCC. Furthermore, the search for genes associated with these variations in cellular composition allows us to identify immunoglobulin genes linked to ICB response. We also develop the Tumor-Immunity Differential (TID) score, based on the tumor mutational burden together with the cell fractions in metastatic ccRCC samples, which is highly predictive of ICB response. Finally, among the TID-related genes, we identify *YWHAE* as an accurate and robust predictor of ICB response in ccRCC, melanoma and non-small cell lung cancer (NSCLC). Overall, we demonstrate that variations in TME composition estimated by cell deconvolution from ccRCC metastatic sites and the gene signatures derived therefrom are highly predictive of patient response to ICB therapies.

## RESULTS

### Comprehensive identification of cell types in ccRCC tumor micro-environment from single-cell RNA seq data

A comprehensive literature review was performed to identify markers, mainly cell surface proteins, known to be specific to certain cell types present in the tumor micro-environment. This led us to construct a hierarchical decision tree summarizing the state of the art associations between cell surface markers and the identity of different cell types present in ccRCC and its tumor micro-environment (fig. S2). Single-cell RNA-seq data generated from 11 treatment-naive resected ccRCC tumors were then collected from a previously published study In this study, immune- and non-immune cells had been separated by CD45-based fluorescence-activated cell sorting (FACS), which we used to separately analyze CD45+ and CD45-cells. We completely reprocessed this dataset through pre-processing, quality control, normalization and cell clustering steps. We then relied on our hierarchical decision tree of cell surface markers to identify the different cell types present in this data set. Based on our reanalysis of the cells retrieved from all patients, we could define 21 distinct cell types, including 3 tumor cell types (Tumor cells, CA9 High Tumor cells, Cycling Tumor cells) (Figs. 1A, B). We assessed these identifications by controlling gene expression of all the specific gene markers of immune and non-immune cells (figs. S3, S4). In addition, the tumor cell populations were examined by the prediction of copy number variants (CNV) (fig. S5). We observed that annotated tumor cell types harbored a deletion of chromosome 3p region (VHL, PBRM1, BAP1, SET2D) and an amplification of the 5q region, which is common for ccRCC (*35, 36*).

**Fig. 1.**
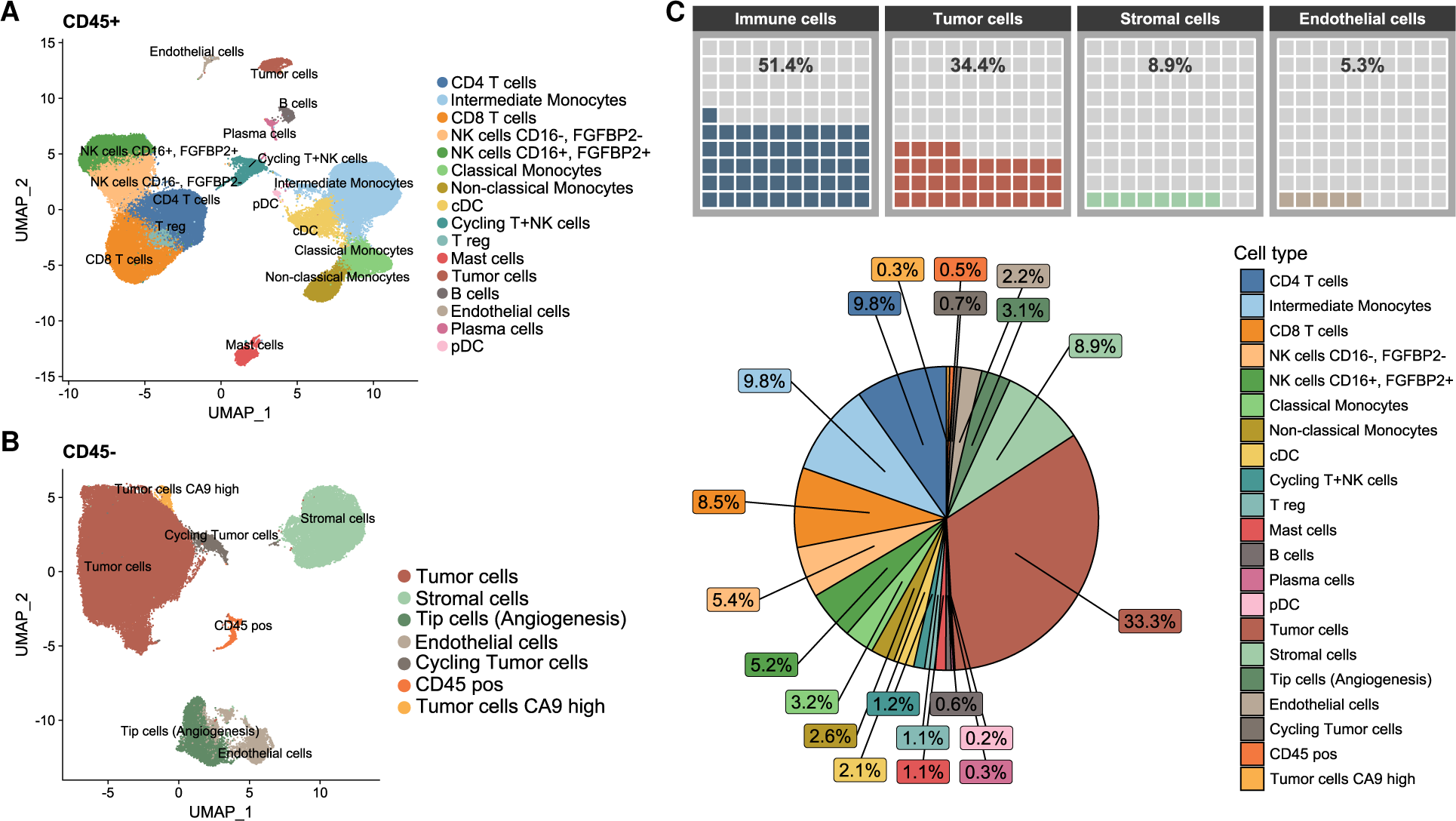
Extended annotation of ccRCC microenvironment cell types from single-cell RNA-Seq from Obradovic et al. (**A)** UMAP of the CD45-positive cells. (B) UMAP of the CD45-negative cells. (C) Distribution of the cell type proportions. The cell fraction values in the pie chart can be read counterclockwise from a cell type compared to the legend cell list (e.g, Stromal cells are following the Tumor cells counterclockwise in the pie chart).

When pooling the cells from all patients, we showed that 34.4% were tumor cell types, 8.9% were stromal cells, 5.3% were endothelial cells, and 51.4% were immune cell populations (Fig. 1C). The latter was mainly composed of monocytes (15.6 %), NK cells (10.6%), CD4-T (9.8%) and CD8+ T (8.5%). These results are consistent with other single-cell analyses of immune cell proportions from ccRCC tissues (*37*).

### Cell deconvolution provides a robust prediction of the cell type composition of ccRCC TME

Transcriptome-based cell deconvolution approaches make it possible to predict the respective proportions of cell types present in a tissue from its bulk transcriptome (*38*). Building on our extensive identification of ccRCC tumor cells and other cell populations defining the TME, we assessed the relevance of using these single-cell RNA-seq data as reference profiles for cell deconvolution of bulk RNA-seq data. We selected the CIBERSORTx and MuSiC methods (*39, 40*), considered by recent benchmarks to be among the most accurate (*38*), and evaluated their ability to predict the respective proportions of the 21 cell types referenced in our single-cell data. To perform this evaluation, we generated test datasets corresponding to 100 pseudo-bulk mixtures by sampling and mixing in controlled and known proportions of the reference single-cell RNA-seq profiles (data file S1). In this way we avoided technical or biological biases between our test datasets and the reference single-cell data. We predicted the proportions of cell types for each pseudo-bulk mixture (data file S2) and assessed the robustness of the estimated fractions obtained by cell deconvolution by comparing them to the expected cell type fractions. We found high correlation values (Rho ranging from 0.57 to 0.99, two-sided Spearman correlation test p-value < 0.0001) between predicted and expected cell type proportions associated with low RMSE (Root Mean Squared Error) dispersion values (Fig. 2A). These predictions were also associated with a very low level of cross-correlation between different cell types, illustrating the specificity of the marker genes used to label single-cell RNA-seq data and the ability of these cell types to be interrogated by cell deconvolution (fig. S6A). In addition, the analysis of the slope of the regression line by cell type between predicted and expected proportions showed good linearity often close to 1, illustrating the quantitative accuracy of the predicted cell type fractions (Fig. 2A, fig. S6B). A comparison of the quality of the predictions generated by CIBERTSORTx with those of the MuSIC method, also widely used in cell deconvolution, was performed. Evaluation of cell proportions produced by MuSIC using pseudobulk mixtures showed that they were of good quality but with lower correlation values and regression coefficients compared to those obtained with CIBERTSORTx (figs. S7A, S7B). In order to complete the quality assessment of the cell fractions predicted by deconvolution by cell type across the patients, an evaluation was carried out this time by patient across the cell types (fig. S7C). We found that MuSIC predictions were strongly correlated with pseudobulk mixtures, but still at a lower level than those produced by CIBERSORTx. This led us to continue further assessment of cell deconvolution using CIBERSORTx only.

**Fig. 2.**
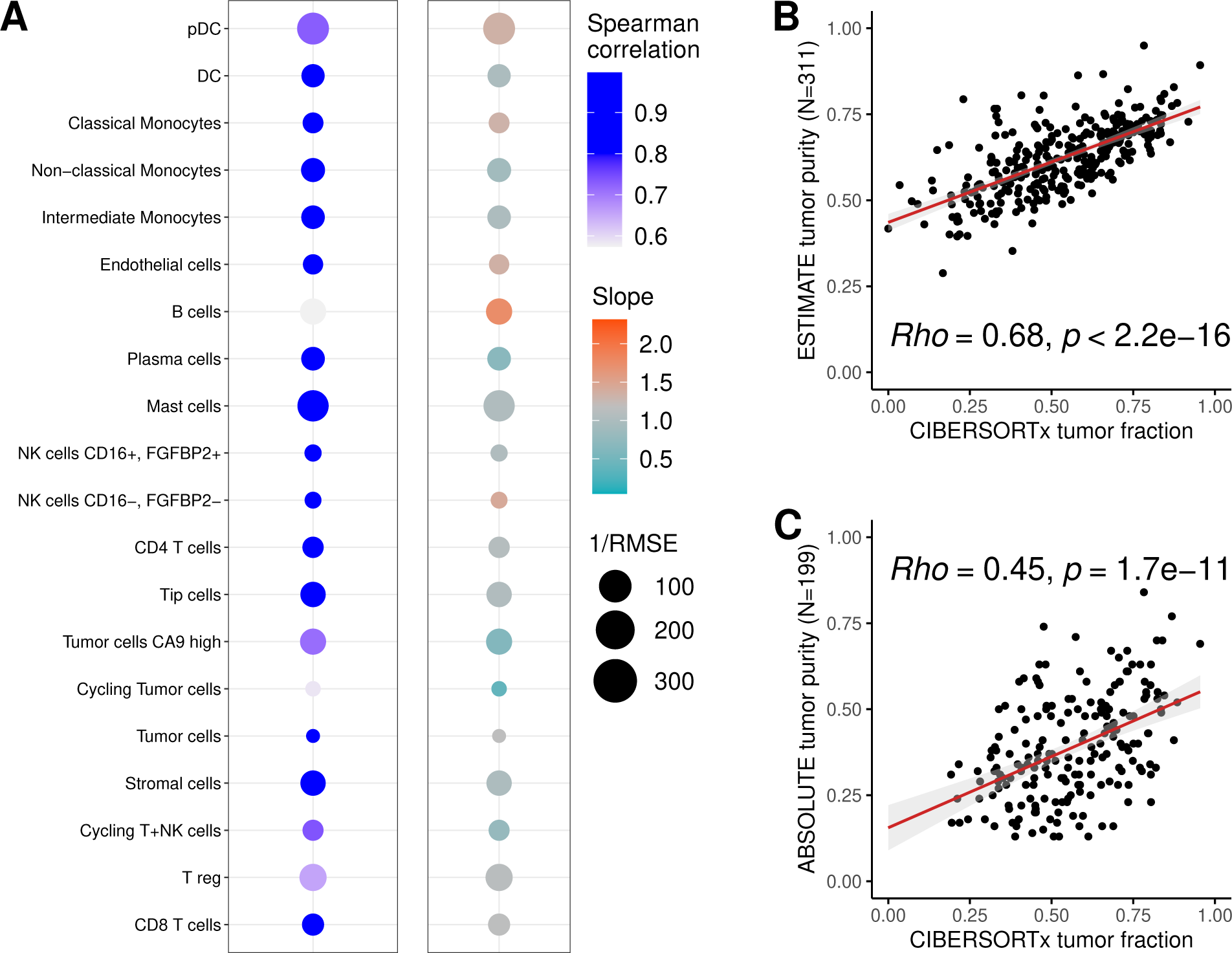
Computational assessment of cell fractions estimated by cell deconvolution. (**A**) Correlation levels between simulated pseudo bulk single-cell RNA-seq mixtures with known proportions and fractions predicted by cell deconvolution using CIBERSORTx. Correlation values are assessed using Spearman correlation coefficients, slopes of the best linear regression line and inverse of the RMSE values. (B) Correlation between tumor purity values calculated by ESTIMATE method and tumor fractions estimated by CIBERSORTx. (C) Correlation between tumor purity values calculated by ABSOLUTE method and tumor fractions estimated by CIBERSORTx.

### Tumor purity of ccRCC is reliably estimated by cell deconvolution

Prediction of tumor purity of tissues from genomic or transcriptomic data is often performed using the ABSOLUTE (*41*) and ESTIMATE (*42*) algorithms, respectively. Here we wanted to explore and assess the ability to predict the tumor purity using a cell deconvolution method based on our reference single-cell RNA-seq profiles. To our knowledge, this possibility has not yet been explored on ccRCC data. We compared tumor purity levels predicted by these different approaches using RNA-seq data from a previously published cohort of 311 ccRCCs (*2*). First, we predicted by cell deconvolution with CIBERSORTx the proportions of the 21 cell types for each of the 311 transcriptomes. For the calculation of tumor purity, we summed the proportions of the 3 tumor cell types. Besides, we used the ESTIMATE tool on these same 311 transcriptomes to predict a second tumor purity value. Finally, we collected from the supplementary data associated with the cohort a third tumor purity value generated by the ABSOLUTE tool. We observed that the deconvolution-based estimated fractions of tumor cells were highly correlated to values predicted by ESTIMATE (two-sided Spearman correlation test p-value < 2.2e-16) (Fig. 2B) and ABSOLUTE (two-sided Spearman correlation test p-value=1.7e-11) (Fig. 2C). We next extended this assessment of tumor cell fractions predicted by cell deconvolution for two other datasets. We generated predictions of tumor cell fractions on a pseudobulk mixture generated from single-cell RNA-seq data from 7 ccRCC samples (*43*) and obtained results very close to those directly quantified from single-cell RNA-seq (fig. S8A). We also predicted the fractions of tumor cells from RNA-seq data of the TCGA ccRCC cohort (*44*) and found that the tumor proportions were significantly correlated and proportional to those generated with the ESTIMATE method (fig. S8B). To conclude, the tumor purity proportions predicted by ESTIMATE and by the cell deconvolution method were consistent, illustrating the relevance of our new approach.

### Cell fractions predicted by deconvolution are consistent with immunofluorescence quantification

We next extended our validation study of cell type fractions estimated by cell deconvolution using cell counts measured by immunofluorescence (IF) for a cohort of 19 ccRCC primary tissues. The ccRCC tissue cohort was profiled by Bulk RNA Barcoding (BRB)-seq to quantify gene expression and imaged by IF to directly enumerate populations of CD8-positive (CD8+), endothelial cells expressing CD34 (CD34+) and CD45-positive (CD45+) leukocytes in red (left panel), yellow (center panel) and green (right panel), respectively (Fig. 3A, data file S3). The bulk transcriptomic data of each sample was analyzed by cell deconvolution with CIBERTSORTx using the support of our single-cell reference data defining 21 cell types. To allow comparison between the cell counts obtained by IHC or IF using the surface markers CD8, CD34 and CD45, we grouped the cell types predicted by cell deconvolution according to the expected presence of these markers in the single-cell data set. Thus, the CD8+ T cells, cycling T+NK cells and NK cells CD16+/FGFBP2+ were considered as CD8+ cells, the endothelial and Tip cells were selected as CD34+ cells and the DC, pDC, classical monocytes, intermediate monocytes, non-classical monocytes, B cells, plasma B cells, cycling T+NK, NK cells CD16-/FGFBP2-, NK cells CD16+/FGFBP2+, CD45-pos, T regulatory (T reg), CD4-T, CD8+ T and mast cells were grouped as CD45+ cells. For each of the 19 samples we summed the predicted cell fractions as described above, to estimate the proportions of CD8+, CD34+ and CD45+ cells and compare these estimations to the IHC or IF measurements. We also extracted the respective contributions, predicted by cell deconvolution, of the cell types composing the 3 populations. As expected, the CD8+ population was predicted to be predominantly composed of the CD8+ T cell type (Fig. 3B left panel). The CD34+ population seemed to be constituted in a balanced manner by endothelial and Tip cells (Fig. 3B center panel). As for the CD45+ population, cell deconvolution predictions seemed to reveal a composition in favor of DC, intermediate monocytes, B and NK cells (Fig. 3B right panel). We observed significant correlations between IHC-based quantification and CIBERSORTx-based estimation for the CD8+ and CD34+ cells (two-sided Spearman correlation test P < 0.05) (Fig. 3C left and center panels) and a positive trend for the CD45+ cells (two-sided Spearman correlation test P = 0.075) (Fig. 3C right panel). To conclude, these analyses demonstrated the reliability of our cell deconvolution approach to predict cell type composition of ccRCC samples.

**Fig. 3.**
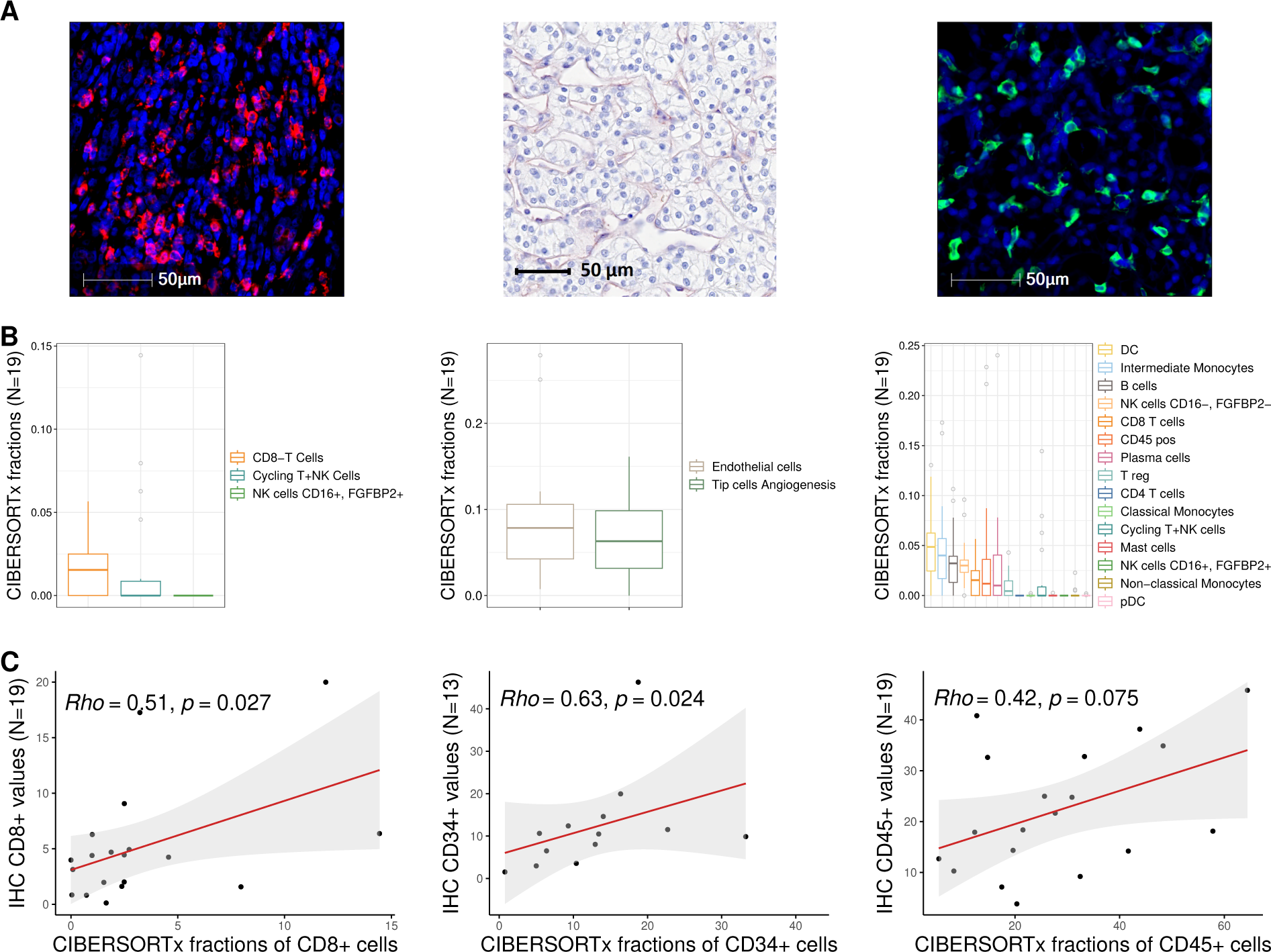
Experimental evaluation of cell fractions estimated by cell deconvolution. (**A)** Examples of single plex immunofluorescence (IF) or immuno-histochemistry (IHC) for CD8+, CD34+, and CD45+ cells (left panel: red for CD8, center panel: red for CD34, right panel: green for CD45), Hoechst counterstaining was used to color nuclei. (B) Distribution of estimated cell fractions of CD8+, CD34+ and CD45+ cells for ccRCC tissue samples by cell deconvolution. (C) Correlation between IHC/IF-based and CIBERSORTx-based quantification of CD8+, CD34+ and CD45+ cells from 19, 13 and 19 matched tumors, respectively.

### The micro-environment of ccRCC metastases predicts patient response to Nivolumab

The tumor microenvironment is considered to be a key factor to understand and predict the response of patients to anti-tumor treatments, in particular to immunotherapies (*25, 26, 29, 30*). The cell deconvolution approach offers the possibility of analyzing the transcriptomic data of large tumor tissue cohorts, of patients included in clinical drug trials, in order to reveal the composition of their TME. In the previous section, we validated our cell deconvolution approach based on the CIBERTSORTx algorithm and single-cell reference transcriptomes of 21 cell types of the ccRCC microenvironment. In a next step we used this approach to predict by deconvolution the cell fractions for each transcriptome of the 311 tumor samples from patients included in the clinical trials CheckMate of the anti-PD-1 antibody Nivolumab (anti-PD-1) or the mTOR inhibitor Everolimus in advanced ccRCCs (CM-009 (NCT01358721), CM-010 (NCT01354431) and CM-025 (NCT01668784)) (*2*). This cohort was selected because it was one of the few to be composed of samples obtained from both the primary tumors and from metastases, associated with clinical metadata describing their responses to therapy (Clinical Benefit (CB), Progression Free Survival (PFS), Overall Survival (OS)).

First, to control a putative sex-related effect, we estimated cell fractions for primary and metastatic tissues and did not find a significant difference in TME composition between men and women after correction for multiple testing (figs. S9, S10). Next, regardless the gender, we subdivided the cohort into 4 subgroups based on sample collection site (primary/metastasis) and therapy (Everolimus/Nivolumab). The cell type proportions were then predicted by cell deconvolution for each subgroup. We observed no statistical association between variations in TME and patient response to Everolimus from the subgroup of 92 primary tissues or of 37 metastases (figs. S11-14, data files S4, S5). We similarly studied the subgroup of 133 primary tumor samples from patients treated with Nivolumab and again did not observe a significant association between the TME composition and clinical variables of response to treatment or survival (figs. S15, S16, data file S6).

We finally selected, among the 311 ccRCC samples of the overallcohort, the 84 samples derived from metastases, and further reduced our selection to the 47 samples taken before the treatment of patients with Nivolumab (Fig. 4A). We performed statistical association analyses between the cell fractions of each of the 19 cell types, and the three categories of patient clinical benefit (Clinical Benefit (CB), Intermediate Clinical Benefit (ICB), Non-Clinical Benefit (NCB) defined by Braun et al. (*2*). Predicted cell fractions of Tumor, T regulatory, CD8+ T and Plasma cells were significantly associated with CB or NCB responders before multiple testing procedure (two-sided Wilcoxon rank-sum test: p-value < 0.05, Fig. 4B). These 4 cell types identified as key populations of the ccRCC TME were kept for further analysis.

**Fig. 4.**
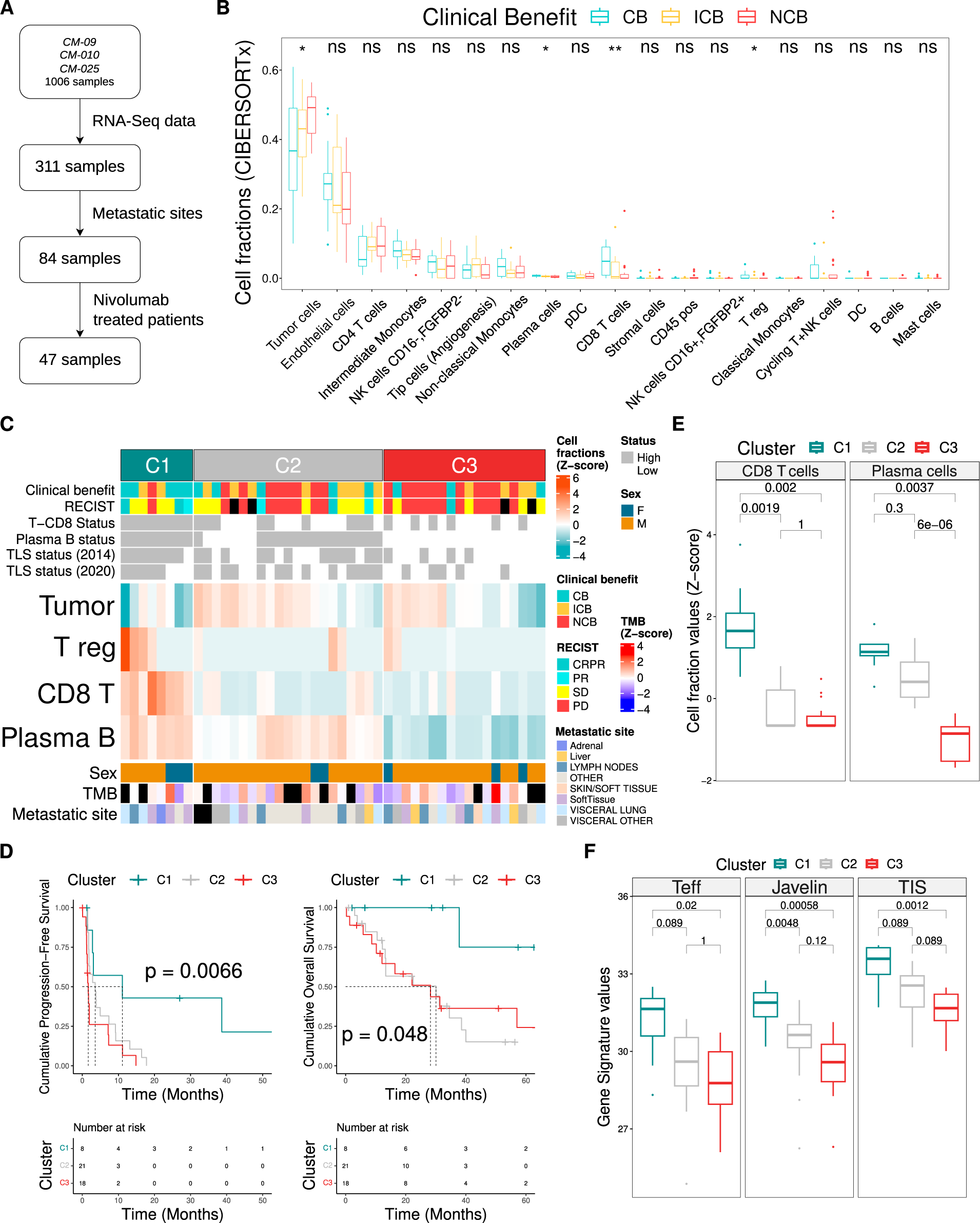
TME subtypes of ccRCC metastases related to ICB treatment response. (**A**) Selection of pre-treatment metastatic site samples from patients treated with nivolumab in the CheckMate cohort. (B) Measures of cell fractions estimated by CIBERSORTx according to ICB clinical benefit. (C) Heatmap of the unsupervised consensus clustering of the metastatic site samples into 3 TME subtypes based on estimated cell fractions (black rectangles represent missing values). Cell fraction status was determined for a given cell fraction by its median value to divide samples into High or Low clusters). (D) PFS and OS analyses based on TME subtypes. (E) Measures of cell fractions of CD8 T cells and plasma cells according to TME subtypes (F) Published gene signature values according to TME subtypes.

### C1-C3 subtypes are associated with tertiary lymphoid structures and ccRCC recurrence to Nivolumab

In order to reveal cell fraction profiles associated with patient clinical outcome to Nivolumab, we performed a consensus clustering of k-means of the 47 metastatic ccRCC samples, based on Tumor, T regulatory, CD8+ T and Plasma cell fractions, and classified them into three subtypes (C1-C3) (Fig. 4C, fig. S17). Among these subtypes, tumors in C1 showed better treatment response outcomes than tumors in C3 (two-sided chi-squared test p-value < 0.05), the latter being characterized by a high proportion of poor response cases (NCB). The C2 subtype groups together a population of patients with varying clinical benefit. We found that the tumor shrinkage values, which reflects the change in size of the tumor targets after treatment, were significantly different between C1, C2 and C3 subtypes with mean values of −34, −6.4 and +16, respectively (Kruskal-Wallis test, p-value < 0.05) (data file S7). We finally performed survival analyses and found that ccRCC tumors from C1 subtype harbored greater PFS (two-sided log-rank test p-value = 0.0066) indicating less post-treatment progression and greater OS (two-sided log-rank test p-value < 0.05) than C2-C3 subtypes (Fig. 4D). As for metastatic sites, TMB counts, gene mutation profiles, tumor purity, sex and age were not different between these 3 subtypes (Kruskal-Wallis or two-sided chi-squared tests, p-value > 0.05, data file S8)

Regarding the TME content, the C1 subtype was associated with a higher fraction of CD8+ T cells and of plasma cells compared to C2-C3 and C3, respectively (Fig. 4E). Plasma cell fractions were able to distinguish the mixed C2 subtype from the C3 subtype of poor responders, the latter being associated with significantly lower fractions (Fig. 4C, E). Tertiary Lymphoid Structures (TLS) have been highlighted as key actors for sustaining an immune-responsive micro-environment (*29, 32, 45*), hence, we analyzed the TLS status in our subtypes. Two TLS scores were calculated using Dieu-Nosjean et al. and Cabrita et al. gene signatures (*29, 45*). The median of each TLS score was used to assign TLS-High and TLS-Low status to the 47 metastatic ccRCC samples. We observed that the C1 subtype was enriched in TLS-High status compared to the C3 subtype (TLS2014: two-sided chi-squared test, p-value < 0.01) (Fig. 4C, data file S8).

To further characterize the mechanisms contributing to the differences of patient response outcomes between C1, C2 and C3 subtypes, we used the following gene signatures calculated in Braun et al.: angiogenesis (Angio) (*46*), myeloid cell infiltration (Myeloid) (*46*), T effector cell infiltration (Teff) (*46*), Tumor Inflammation Score (TIS) (*47*), and immune infiltration (JAVELIN) (*1, 2*). We observed that C1 subtype tumors were associated with greater Teff, JAVELIN and TIS scores, compared to C3, C2-C3 and C3, respectively (two-sided Wilcoxon rank-sum test, Benjamini-Hochberg-corrected p-value < 0.05, p-value < 0.01 and p-value < 0.01) (Fig. 4F). These results confirmed the high immune infiltration, in particular of T cells, and the inflammation of C1 subtype samples. These biological features are known to promote the beneficial response of patients to immunotherapies (*1, 46, 47*), which is coherent with cluster C1 being enriched in patients with a clinical benefit.

### Immunoglobulin-related gene expression values are associated with the C1-C3 clusters and patient response to Nivolumab

We next aimed at investigating gene expression values associated with the three subtypes (C1-C3). We leveraged the reprocessed and relabeled single-cell RNA-seq data of the ccRCC and performed pair-wise comparisons between cell-type transcriptomes to identify sets of differentially expressed genes specific to each cell type (Fig. 5A, data files S9, S10). To do so, only the gene sets of the 4 cell types previously identified in metastases to be associated with the response of patients to Nivolumab (Tumor, T regulatory, CD8+ T and Plasma B cells, Figs. 4B and C) were considered. Finally, we reduced the list of genes to those being significantly differentially expressed between the C1, C2 or C3 cell-based subtypes (two-sided Wilcoxon rank-sum test, Benjamini-Hochberg-corrected p-value < 0.05) (Fig. 5B). This gene selection process revealed 6 genes: *TMEM139*, whose reduced expression is associated with a poor prognosis in Non-Small Cell Lung Cancer (NSCLC) (*48*) and 5 immunoglobulin genes coding for the constant region of immunoglobulin light chains (*IGKC*) and heavy chains (*IGHG1*, *IGHG2*, *IGHG3*, *IGHA1*). As expected, the analysis of these 6 genes by functional enrichment of biological processes (GO) indicated its involvement in the tumor immune response mediated by B cells (fig. S18, data file S11).

**Fig. 5.**
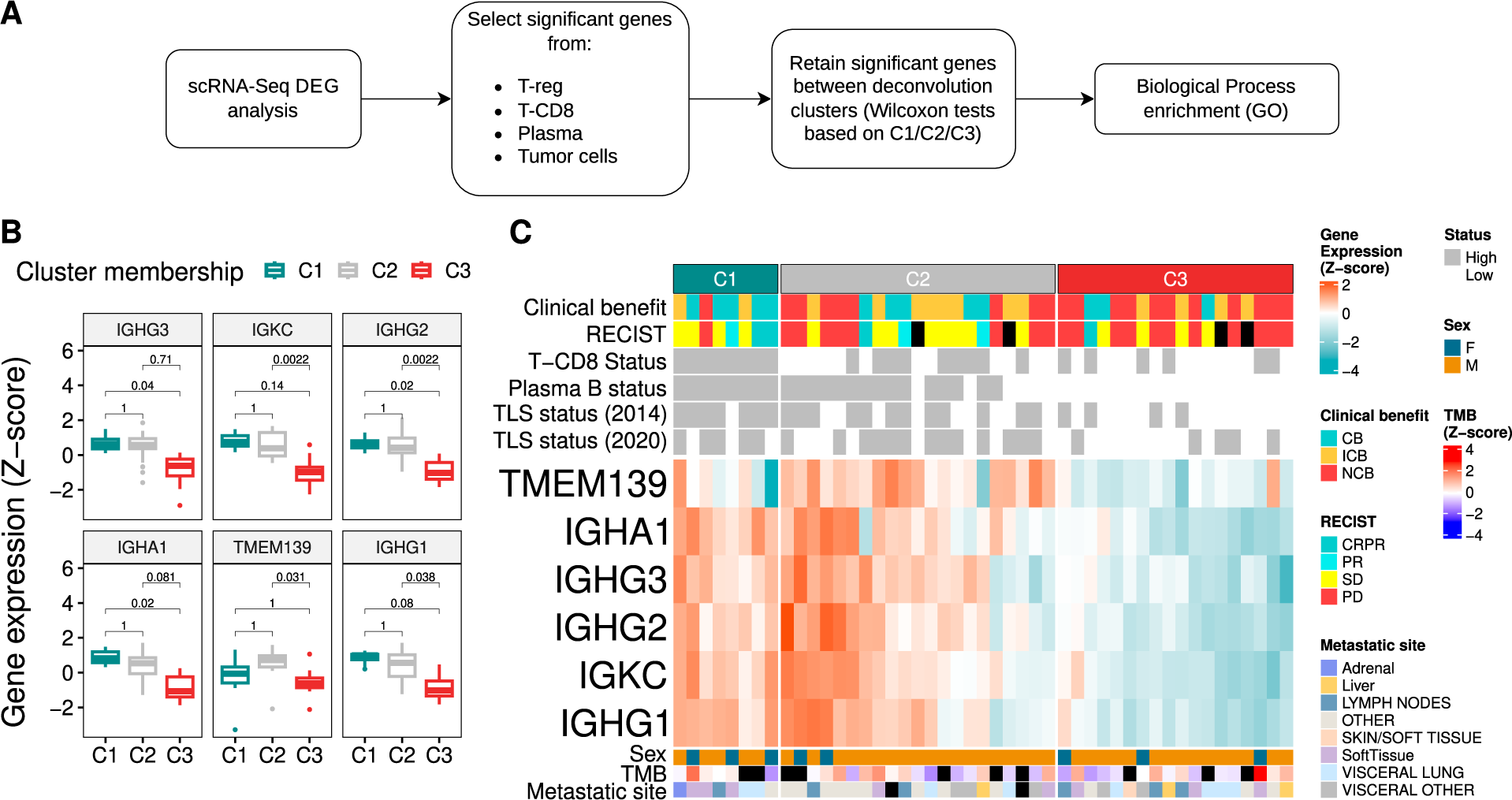
Gene expression values associated with TME subtypes. (**A**) Selection of genes according to single-cell RNA-Seq DEG analysis and the TME-subtypes. (B) Measures of the gene expressions significantly associated with TME subtypes. (C) Heatmap of gene expression values and cell fraction status (a given cell fraction was divided by its median values to give High or Low clusters) according to the TME-subtypes (black rectangles represent missing values).

We observed that the immunoglobulin gene expression values were higher for the C1 cluster compared to C3 (Fig. 5B). This was consistent with the fact that tumors from C1 were characterized by a higher proportion of plasma B cells, which are immunoglobulin-secreting cells, compared to C3 (Figs. 4E, Fig. 5C). The C2 subtype also had stronger immunoglobulin gene expression values compared to C3. However, the heterogeneity of treatment responses in this C2 group could be due to the lower fraction of CD8+ T cells compared to the C1 subtype of good responders. To conclude, the expression of immunoglobulin genes combined with the proportion of plasma and CD8+ T cells in ccRCC metastases would be key determinants in predicting patient response to Nivolumab.

### Tumor-Immunity Differential score predicts patient response to Nivolumab from ccRCC metastases

Leveraging on our results, we found that a wide variety of molecular and cellular properties of ccRCC metastases help subdivide patients into profiles associated with response to Nivolumab. In order to integrate these components, we developed a Tumor-Immunity Differential (TID) score. The TID score consists of the difference between a tumor part of high predictive value for patients with NCB (Tumor cell fraction and TMB) and an immunity part of high predictive value for CB patients (Plasma B, CD8+ T and Treg cell fractions and the *PDCD1* gene expression) (Figs. 6A and 6B).Therefore, the higher the TID score the worse the patient’s prognosis should be. To assess this hypothesis, the cohort of 47 ccRCC metastases was divided using the TID score median value (0.78) resulting in 24 tumors with TID-Low status and 23 tumors with TID-High status.

**Fig. 6.**
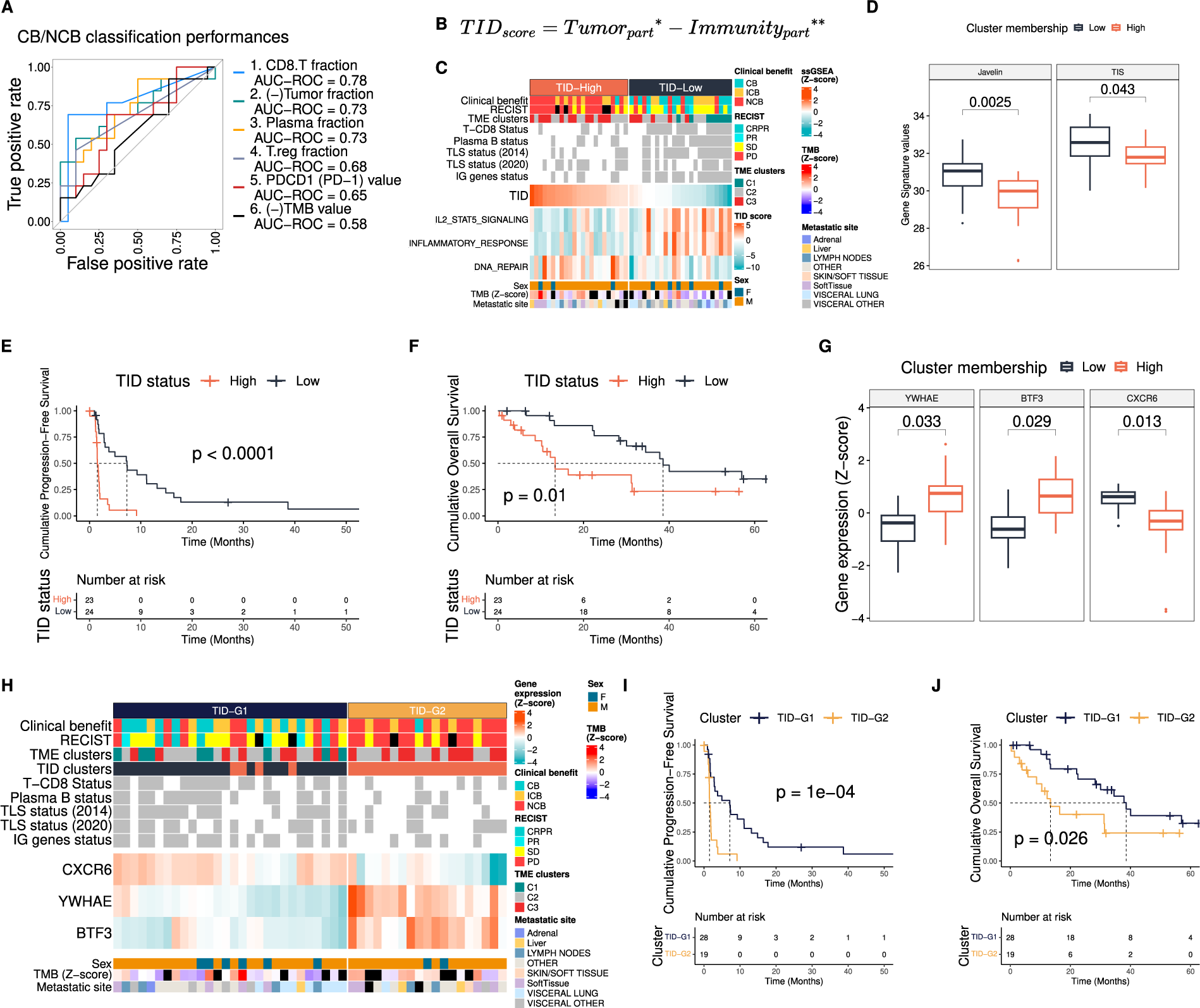
Tumor-Immunity Differential (TID) score associated with ICB treatment response. (**A**) ROC curves and AUC-ROC values for estimated cell fractions and gene expressions related to ICB treatment response. (B) TID score formula (**\***The tumor part is the sum of the tumor fraction and the TMB values, **\*\***The immunity part is the sum of the T-CD8, Plasma B and T regulatory fractions and the *PDCD1* gene values). (C) Clustering of metastatic site samples based on the TID score divided into TID-High and TID-Low by its median value (black rectangles represent missing values). Cell fraction status was determined for a given cell fraction by its median value to divide samples into High or Low clusters). (D) Published gene signature values according to TID subtypes (E) PFS and (F) OS values of the TID subtypes. (G) Expression values of differentially expressed genes between TID subtypes. (H) Unsupervised consensus clustering of the metastatic site samples into 2 clusters, TID-G1 and TID-G2 subtypes, based on the 3 genes related to TID subtypes (*CXCR6*, *YWHAE*, *BTF3*) (black rectangles represent missing values). (I) PFS and (J) OS values of the TID-G1 and TID-G2 clusters.

To characterize these groups, we performed a ssGSEA analysis of biological hallmarks from the MSigDB database. Three gene sets yielded significant differences between TID-subtypes. The TID-High profile was characterized by a lower enrichment of the immunity-related hallmarks, INFLAMMATORY_RESPONSE and IL2_STAT5_SIGNALING, and by a higher enrichment of the DNA_REPAIR process (Fig. 6C) (data file S12).

Furthermore, we showed that the TID-High subtype had significantly worse clinical prognostic values and was enriched in patients with NCB responses for 15 out of 23 tumors (the 8 left consisting of 1 CB and 7 ICB) whereas the TID-Low subtype was enriched in patients with CB responses for 12 out of 24 tumors (the 12 others being 5 NCB and 7 ICB) (two-sided chi-squared test, p-value < 0.001) (Fig. 6C). Furthermore, the TID subtypes were significantly associated with patient PFS and OS (two-sided log-rank test, PFS: p-value < 1e−04 and OS: p-value = 0.01) (Figs. 6E and 6F). The TID-Low subtype of good prognostic also harbored lower tumor purity (ABSOLUTE value 0.28 vs. 0.39, data file S13) and higher JAVELIN and TIS scores compared to TID-High subtype (two-sided Wilcoxon rank-sum test, Benjamini-Hochberg-corrected p-value < 0.05), highlighting for the latter a higher tumor content associated with a reduced tumor immune infiltrate (Fig. 6D).

We continued our investigation of TID subtypes by identifying genes whose expression significantly differed between these two groups. Only three genes were found to be significantly differentially expressed between the two TID subtypes (two-sided Wilcoxon rank-sum test, Benjamini-Hochberg-corrected p-value < 0.05) (Fig. 6G). The *CXCR6* gene was under-expressed in TID-Low cluster, while the *YWHAE* and *BTF3* genes were over-expressed in TID-High subtype.

Since the TID score integrates several molecular and cellular components that can be challenging to obtain in clinical routine, we studied the ability of this 3-gene signature to classify the cohort of 47 ccRCC metastases, using consensus clustering based on k-means, into TID-associated clusters TID-G1 and TID-G2 (Fig. 6H). These clusters were significantly associated with PFS, OS values and RECIST treatment outcomes with the TID-G1 showing a better treatment outcome and OS than TID-G2 (two-sided log-rank test, PFS: p-value = 1e−04, OS: p-value < 0.05) (Figs. 6I and 6J, data file S14). Interestingly, *CXCR6* showed a higher expression in TID-G1, compared to TID-G2, and associated with higher proportion of CD8+ T and plasma B cells, higher expression of immunoglobulin genes, and higher TLS scores (Fig. 6H). These results demonstrated that the 3-gene signature was able to capture the treatment response discrepancy reflected by the TID score from ccRCC metastases.

### TID-associated *YWHAE* gene predicts patient response to Immune Checkpoint Blockade in melanoma and lung cancers

To explore the predictive power of the TID score and TID-associated genes (*CXCR6*, *YWHAE* and *BTF3*), we performed a comprehensive comparison to previously published Immune Checkpoint Blockade (ICB) response predictive scores. We used independent gene expression datasets based on primary and metastatic tissues from various cancers from patients treated by anti-PD-1 or anti-CTLA-4 therapies. The ICB-related scores considered were transcriptome-based predictive signatures, including JAVELIN (*1*), MIAS (*22*), GEP (*20*), MHC-I (*21*), MHC-II (*21*), IMPRES (*19*) two TLS scores (TLS2014 (*29*) and TLS2020 (*45*)), and the PD-1 gene (*PDCD1*) expression level.

We initiated our comparison using three cohorts of ccRCC tissue obtained from patients before treatment with anti-PD-1: a first of 33 metastases (*2*), a second of 90 primary tumors (*2*) and a third of 9 primary tumors (*49*) (Fig. 7A). We found that the TID score was the best performing predictor of anti-PD-1 response in ccRCC metastases (AUC-ROC of 0.88), followed by MHCI and MHCII scores (AUC-ROC of 0.8 and 0.78) (Fig. 7A). We also observed good performances for the three TID-associated genes (AUC-ROCs ranging from 0.73 to 0.76). Surprisingly, all the ICB response scores calculated with primary ccRCC samples from the Braun et al. dataset obtained poor performances close to chance (AUC-ROCs ranging from 0.5 to 0.61) while the mixed dataset of primary and metastatic samples from Ascierto et al. harbored slightly better performances, especially for the *YWHAE* gene with an AUC-ROC of 0.75 (Fig. 7A, data files S15-17). To explain the discrepancy between the score performances for the ccRCC datasets, we assessed the correlations between both types of samples, primary and metastatic. We used an independent cohort of 15 matched primary and metastatic ccRCC samples (*50*) and calculated the same scores based on RNA gene expression data (data file S18, S19). We observed correlations between primary-based and metastatic-based scores only for IG, IMPRES and TLS2020 scores (two-sided Spearman correlation test p-value < 0.05) (Fig. 7B) suggesting that treatment response prediction based on gene expression in ccRCC is not correlated between primary and metastasis samples from the same patient.

**Fig. 7.**
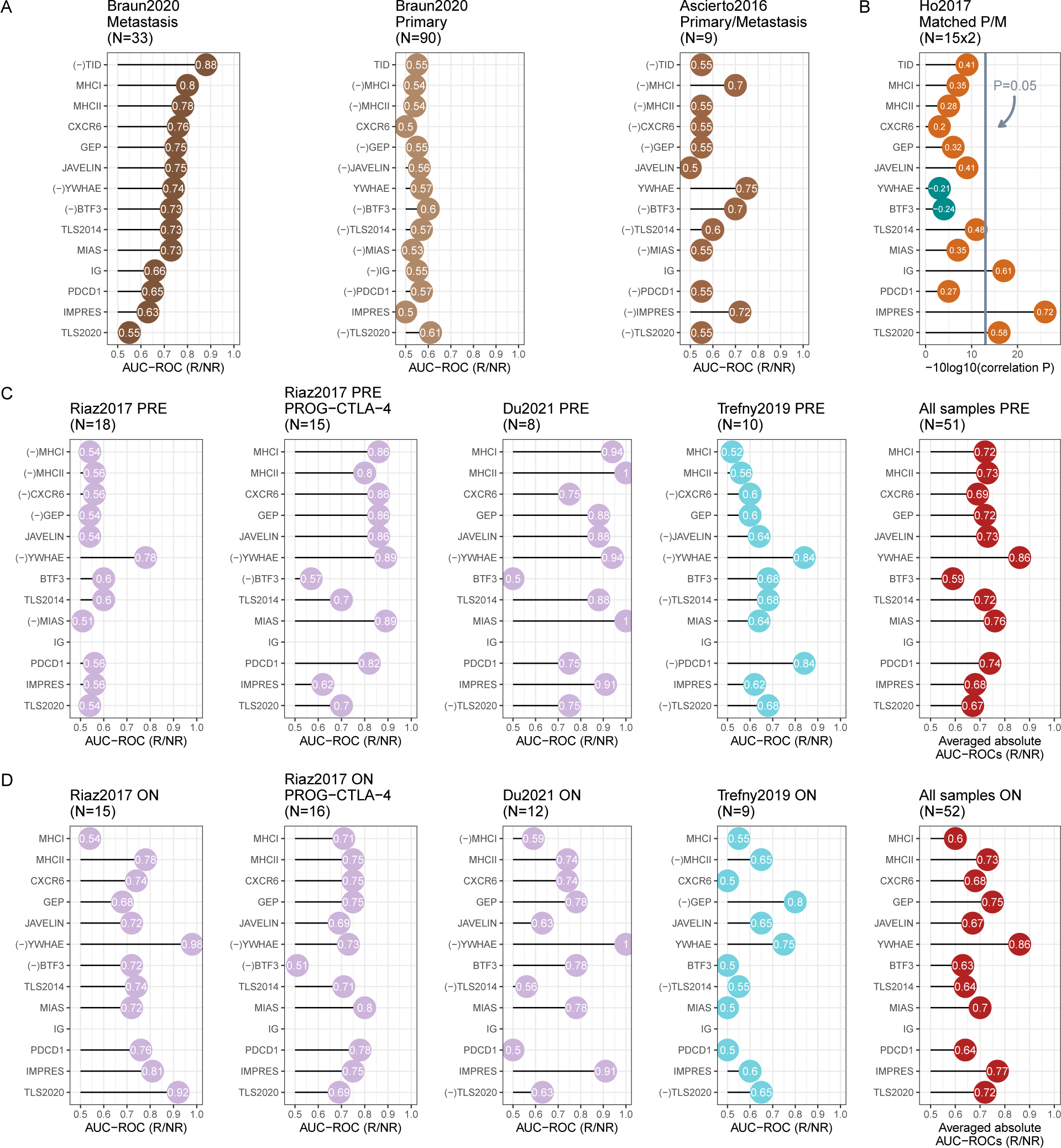
TID score and TID-related genes as key markers of ICB treatment response in several cancer types. (**A**) Classification performances of responders (R) over non-responders (NR) samples for a collection of predictive scores and genes calculated from cohorts of pre-treated (before Nivolumab treatment) ccRCC samples. (B) Correlation between scores calculated for 15 patients with matched ccRCC primary and metastatic site samples. Classification performances of R over NR for (C) pre-treated melanoma and NSCLC (PBMCs) samples and (D) on-treatment melanoma and NSCLC (PBMCs) samples. (‘Primary’ and “Metastasis’ labels mean that gene expression were obtained from primary or metastatic samples, respectively. ‘Primary/Metastasis’ refers to a mixed dataset of primary and metastatic samples. ‘PRE’ and ‘ON’ labels refer to dataset with pre-treated or on-treatment samples, respectively. ‘PROG-CTLA-4’ label refers to samples who have progressed after an anti-CTLA-4 therapy).

To further investigate the value of TID-associated genes beyond ccRCC, we investigated their ability to predict the response to immunotherapies of patients with metastatic melanoma or metastatic non-small cell lung cancer (NSCLC), compared to published ICB response scores. More specifically, we used two cohorts of patients with metastatic melanoma treated with Nivolumab: a first one of 64 primary samples from Riaz et al. (*51*) and a second one of 20 primary samples from Du et al. (*52*) We also used a cohort from Trefny et al. (*53*) of 19 blood samples with peripheral blood mononuclear cells (PBMCs) collected from patients with metastatic NSCLC treated with Nivolumab. In these three studies, samples were collected before and during treatment of which we took advantage by dividing each cohort according to the time of the biopsy, into pre- and on-treatment samples (Figs. 7C and 7D). Furthermore, to assess how prior therapy could affect patient response to ICB, we divided the Riaz et al. cohort into groups of patients who had previously been treated with Ipilimumab (anti-CTLA-4) (31 samples) and those naive for this treatment (33 samples). Of note, as less than 2 out of 5 IG genes have been detected in melanoma and NSCLC transcriptomic datasets, they could not be used to assess the prediction of the immunoglobulin genes.

Interestingly, for pre-treatment melanoma samples, the TID-associated *YWHAE* gene expression greatly outperformed the existing ICB response scoresfor the Ipilimumab-naive patients (AUC-ROC of 0.78) (Fig. 7C first plot, data file S20). In addition, it was the best performer, together with the MIAS score, for the patients who progressed after Ipilimumab treatment(AUC-ROC of 0.89) (Fig. 7C second plot, data file S21), and was among the top predictors in the Du et al. cohort (AUC-ROC of 0.94) (Fig. 7C third plot, data file S22). Regarding PBMCs samples from the Trefny et al. dataset, the expression levels of *YWHAE* and *PDCD1* genes were the best performers (AUC-ROC of 0.84) (Fig. 7C fourth plot, data file S23). The average AUC-ROC values highlighted the *YWHAE* gene as the best marker of ICB treatment response based on pre-treatment biopsies (AUC-ROC of 0.85) (Fig. 7C fifth plot).

We continued our investigation of patients with melanoma or NSCLC using samples obtained during their treatment with Nivolumab. For the Riaz et al. melanoma cohort, we observed an overall improvement in the performance of the ICB response scores for the Ipilimumab-naive samples compared to those obtained from the biopsies taken before Nivolumab treatment (Figs. 7C and 7D first plots). The best predictive power was obtained by far with the *YWHAE* gene for Ipilimumab-naive samples from the Riaz et al. dataset (AUC-ROC of 0.98) (Fig. 7D first plot, data file S24) and for that of Du et al. (AUC-ROC of 1) (Fig. 7D third plot, data file S25). Regarding the melanoma samples from Riaz et al. taken from patients who progressed after treatment with Ipilimumab, we observed, unlike the Ipilimumab-naive patients, rather an overall reduction in the quality of the predictions with a performance for *YWHAE* (AUC-ROC of 0.73) below that of the MIAS score (AUC-ROC of 0.8) (Fig. 7D second plot, data file S26). Finally, the performances obtained for on-treatment samples in the Trefny et al. dataset were close to those obtained from the samples taken from the patients before treatment (Fig. 7D fourth plot, data file S27). The *YWHAE* gene obtained the second-best prediction (AUC-ROC of 0.75), just after that of the GEP score (AUC-ROC of 0.8), with a strong decrease in the performance of *PDCD1*. Overall, the average AUC-ROCs highlighted the *YWHAE* gene as the best marker of ICB response based on on-treatment biopsies (AUC-ROC of 0.86) (Fig. 7D fifth plot).

## DISCUSSION

Immune checkpoint blockade (ICB) therapies are now an important tool for the treatment of many types of advanced cancers, leading to prolonged progression-free and overall survival(*3, 54, 55*). However, as only a subset of patients responds to ICB therapies, there is an urgent need for novel approaches to better select patients who may benefit from these treatments (*55, 56*). Previous studies have shown the influence of the cellular composition of the tumor microenvironment (TME) on the response of patients to immunotherapies (*31, 32, 45*). Although substantial effort has been devoted to the investigation of T cell populations toward understanding ICB treatment response, other cell types in the TME are also involved in patient clinical outcome (*25, 26*).

As of yet, only limited number of patients have been analyzed by single cell sequencing, hampering identification of response-related aspects of TME composition. In contrast, many clinical trials comprise large series of patients analyzed by bulk RNA-seq. Predicting the composition of the TME from bulk RNA-seq data by cell deconvolution may bridge this challenge and has proven to be a robust and sensitive approach, especially when using a tumor-specific single-cell RNA-seq data as a reference of cell types present in TME (*38*). Also, the quality of single-cell transcriptome labeling used as a reference of these methods has a major impact on the predictive performance in the deconvolution procedure. We therefore performed a detailed characterization of the used single-cell RNA-seq data set to obtain a robust classification of cells, resulting in the annotation of 21 distinct cell types (*34*). We then performed a thorough assessment of the robustness of cell fractions predicted by cell deconvolution using this annotated single-cell dataset. By carrying out simulations of pseudobulk RNA-seq mixtures from single-cell RNA-seq data, we ensured our ability to correctly predict the proportions of the different cell types present in the ccRCC tumor samples. Importantly, we also validated that our cell deconvolution predictions fairly reflected the relative proportions of CD8+, CD34+, CD45+ cell populations directly measured by IHC on tissue sections. The annotated single-cell dataset was thus considered as a robust reference to perform cell deconvolution by CIBERTSORTx on bulk transcriptomic data of advanced ccRCC samples from primary and metastatic sites collected from patients before ICB treatment (*2*).

When exploring the predicted cell type proportions of ccRCC samples from Braun et al., 2020, we found that the tumor, Plasma, CD8+ T and T-regulatory cell fractions in metastatic samples displayed significant differences in relation to anti-PD-1 treatment response, cancer progression and overall survival. Based on this observation, we identified three distinct subtypes (C1-C3). Interestingly, we observed no association between tumor composition and treatment response for samples from primary sites or for primary and metastatic samples treated with an mTOR inhibitor (Everolimus). Moreover, differentially expressed genes between C1-C3 subtypes revealed 5 immunoglobulin genes (*IGKC*, *IGHG1*, *IGHG2*, *IGHG3*, *IGHA1*) as markers of the C3 cluster showing the worst ICB clinical response. This cluster is characterized by low expression of immunoglobulin genes and poor fractions of Plasma cells indicating the key role of Plasma cells in the anti-tumor immune response. This need of both B-cell and T-cell fractions for efficient immunotherapy treatment is consistent with previous works in colorectal (*57*), breast (*58*), NSCLC (*59*), head and neck (*60*), and ovarian (*61*) cancers where higher proportions of T-cells or B-cells in TME were associated with improved patient survival.

Besides, previous studies found that B cells associated with tertiary lymphoid structures (TLS) were involved in adaptive immune responses in inflamed and tumor tissues (*29, 30, 45*). These ectopic lymphoid formations lead to the differentiation of key immune cells: tumor-specific B-cells acting either as antigen-presenting cells or tumor antigen-specific antibody-secreting cells and T-cells (*62, 63*). In recent works, TLS were also associated with ICB treatment responses in ccRCC and melanoma with a focus on TLS-associated B cells and gene signatures (*29, 32, 45*). This is consistent with our results obtained using ccRCC metastases from the CheckMate cohorts (*2*), where high levels of CD8+ T cell and Plasma B-cell fractions were correlated with the two TLS signatures. Furthermore, the subtype C1, enriched in both CD8+ T and Plasma B-cells fractions, was related to a high proportion of ICB good responders. This observation may reflect the important interplay between these cell types in TLS, especially in metastasis sites of patients with ccRCC.

In addition, we developed a single-sample Tumor-Immunity Differential (TID) score to leverage gene expression data and estimated TME cell fractions to cluster metastatic ccRCC samples. We based our score on both favorable (*PDCD1* (PD-1) gene expression, T regulatory, CD8+ T and Plasma cell fractions) and unfavorable (Tumor and TMB) features for ICB response. The TID score was then calculated as the difference between the ICB response unfavorable “tumor” and the favorable “immunity” parts. We observed that the TID-High subtype was strongly correlated with bad responders, recurrence and with a poorer overall survival. Also, high TMB values appeared to be associated with bad ICB response in metastatic ccRCC samples. This observation is directly linked with recent works in ccRCC which have shown that high TMB values were associated with poor survival and immune infiltration (*64*) but with a limited predictive clinical value of ICB response (*10, 11*). Moreover, a high tumor purity was found in the High-TID subtype of bad ICB responders, suggesting that tumor fraction or purity were a key component driving ICB treatment response in metastatic ccRCC samples. Three genes were highlighted as markers of the TID subtypes: *YWHAE*, *CXCR6* and *BTF3*. The *YWHAE* gene belongs to the 14-3-3 protein family and was previously found to be associated with advanced ovarian cancers, and poor patient prognosis mediated by the PI3K/AKT and MAPK pathways (*65*). Interestingly, phenethyl isothiocyanate (PEITC) and fusicoccin molecules are found in the DrugBank database to target 14-3-3 proteins. Previous studies revealed various anti-cancer effects of PEITC molecules (*66*), leading to an inhibition of carcinogen metabolism in smokers with lung cancer (*67*) while Fusicoccin-A induces apoptosis in human cancer cell lines in combination with or after IFN-α treatment (*68, 69*). *CXCR6* is a chemokine receptor overexpressed in tumor-infiltrating lymphocytes involved in the recruitment of T cells into RCC tissue (*70*). The Basic Transcription Factor 3 (*BTF3*) is required for the transcriptional initiation and known to be an oncogene in colorectal cancer (*71*). Furthermore, it is overexpressed in pancreatic ductal carcinoma cells (*72*) and in prostate cancer where it sustains a cancer stem-like phenotype (*73, 74*). To further assess the clinical relevance of the TID score and the associated genes, we compared them to existing transcriptomic scores previously published to predict patient clinical response to ICB therapy. We observed that the TID score was the best predictor of patient clinical benefit on the cohort of metastatic ccRCC samples from the CheckMate cohorts (*2*).

In order to assess the predictive power of marker genes associated with the TID score, we extended the comparisons of scores and markers predicting ICB clinical response to melanoma and NSCLC. The classification performances of the TID-associated *YWHAE* gene outperformed other existing scores in anti-PD-1 treated metastatic melanoma and NSCLC cancers in pre-treatment and on-treatment samples. Notably, the best performances in the NSCLC Trefny et al. subgroups were obtained using gene expressions of *YWHAE* and *PDCD1* although the analyzed samples were peripheral blood circulating CD8+ T cells. These results reinforced the clinical relevance of peripheral CD8+ T cells in the prediction of ICB response in NSCLC and may implicate a key role of *YWHAE* in these cells. Interestingly, the investigation of a variety of tumor sample types with different biopsy time-points (pre- or post-treatment by anti-PD-1 antibody) from several cohorts revealed the discrepancy in the prediction performance of treatment outcome by predictive scores based on gene expression data. We observed poorer performances for samples from the primary site of ccRCC compared to the samples from a metastatic site (CheckMate cohorts). In fact, to assess the impact of the sample type, we compared the score values between matched primary and metastatic samples in an independent cohort (Ho et al. dataset). This analysis revealed no significant correlations between the two samples types for good predictors. These results highlighted the critical effect of tumor sample type in the discovery of ICB response predictors.

Also, the prediction performances were not equivalent between pre- and post-treatment samples or samples recruited after the failure of one previous anti-CTLA-4 immunotherapy and without prior treatment in melanoma samples. This is consistent with a previous study on patients with melanoma, treated with Nivolumab or Pembrolizumab, showing that previous anti-CTLA-4 exposure was associated with performance differences for treatment outcome prediction (*21*).

To conclude, we highlighted in this work the importance of the interplay of both CD8+ T and Plasma B cells in immunotherapy response in ccRCC and we revealed the 5 immunoglobulin genes (*IGKC*, *IGHG1*, *IGHG2*, *IGHG3*, *IGHA1*), the Tumor-Immunity Differential score and the TID-associated gene *YWHAE* as powerful markers of ICB treatment response based on pre-treatment or on-treatment biopsies of primary sites, metastatic or peripheral CD8+ T cell samples in several cancer types. Further validation studies in larger cohorts will be needed to assess the ICB predictive performance of the TID score and of the key TID-associated gene markers reported here, both in ccRCC and in other tumor types. Nevertheless, our results open novel avenues to better predict the clinical outcome of patients with cancers treated with immunotherapies.

## MATERIALS AND METHODS

### Public transcriptomics and clinical data

Processed bulk RNA-Seq and clinical data were collected for 5 different cohorts. A cohort of 331 primary and metastatic advanced clear cell Renal Cell Carcinoma (ccRCC) samples from patients that progressed on 1, 2 or 3 previous therapies (at least one systemic anti-angiogenic therapy) included in clinical trials CM-009 (NCT01358721), CM-010 (NCT01354431) and CM-025 (NCT01668784) treated by anti-PD-1 antibody Nivolumab and mTOR inhibitor Everolimus (EGAC00001001519; EGAC00001001520; EGAC00001001521) (*2*). A cohort of 11 primary or metastatic ccRCC samples on one of four clinical trials (NCT00441337, NCT00730639, NCT01354431, NCT01358721) (*49*) processed in (*22*). A cohort of 64 advanced melanoma samples from patients included in clinical trial CA209-038 (NCT01621490) treated with Nivolumab (GSE91061) (*51*). A cohort of 20 metastatic melanoma samples from patients treated by anti-PD-1 (GSE168204) (*52*). A cohort of 19 peripheral CD8+ T cells (PBMCs) samples collected from blood of metastatic NSCLC patients (GSE111414) (*53*). Samples for each cohort were divided according to prior therapies and analyzed independently (fig. S1). Patients with treatment outcomes of “Complete Response” (CR) or Partial Response (PR) were considered as responders, whereas patients with treatment outcome of “Progressive Disease” (PD) as non-responders.

### Single-cell RNA-seq data analysis

Single-cell RNA-seq data from 11 ccRCC patients was obtained from Obradovic et al. (*34*). Raw data was downloaded from https://data.mendeley.com/datasets/nc9bc8dn4m/1. The data set consisted of adjacent normal and tumor tissue. Both were analyzed separately. Furthermore, in the original study, the tissue was prior to sequencing FACS sorted into CD45+ and CD45-subsets. These subset annotations were kept, and the data were separately re-processed in R using Seurat (v.4.1.1) (*75*) and harmony (v.0.1.0) (*76*). Initially, only features expressed in at least 50 cells were included in the analysis. Further QC included removal of cells with fewer than 200 and more than 5000 features detected. As kidney cells have in general a high mitochondrial content, its threshold was set to >25%, everything above was removed. To normalize the data, Seurat’s function NormalizeData was used followed by a feature selection to find the 2000 most variable genes. Afterward, the data was scaled by using the function ScaleData. Prior to data integration, a principal component analysis was carried out. To adjust for patient-to-patient variation, data integration was achieved by the function RunHarmony from the harmony package. For dimensional reduction, uniform manifold approximation (UMAP) was performed on the first 20 dimensions of the harmony reductions. Finally, the Louvain algorithm implemented in Seurat was used for cluster analysis with a resolution of 0.5. To assign cell types to the clusters, differential gene expression (DGE) was used by applying Seurat’s function FindAllMarkers. In addition to DGE, previously known cell type markers were used to facilitate cell type annotation.

To confirm the tumor origin of the annotated tumor cells, copy number variation (CNV) was inferred using the R package infercnv (v.1.12.0) (*77*). As a normal reference the adjacent normal CD45-subset was used. The gene order file (hg38_gencode_v28.txt) was downloaded from https://data.broadinstitute.org/Trinity/CTAT/cnv/. For the analysis a gene cutoff of 0.1 was set. Furthermore, cluster_by_groups was set to TRUE and a noise filter was applied by setting sd_amplifier to 1.5. The hidden Markov model (HMM) i6 was used for CNV prediction.

### Bulk RNA Barcoding (BRB) library preparation and sequencing

Total RNA was extracted from MCTS using the MirVana PARIS kit (Thermofisher). BRB-seq experiments were performed at the Research Institute for Environmental and Occupational Health (Irset, Rennes, France) according to the published protocol (Alpern et al, 2019). Briefly, the reverse transcription and the template switching reactions were performed using 4 µL total RNA at 2.5 ng/µL. RNA were first mixed with 1 µL barcoded oligo-dT (10 µM BU3 primers, Microsynth), 1 μL dNTP (desoxyribonucleoside triphosphate) (0.2 mM) in a PCR (Polymerase Chain Reaction) plate, incubated at 65 °C for 5 min and then put on ice. The first-strand synthesis reactions were performed in 10 µL total volume with 5 µL of RT (Reverse transcription) Buffer and 0.125 µL of Maxima H minus Reverse Transcriptase (Thermofisher Scientific) and 1 µL of 10 μM template switch oligo (TSO, IDT). The plates were then incubated at 42 °C for 90 min and then put on ice.

After reverse transcription (RT), decorated cDNA from multiple samples were pooled together and purified using the DNA Clean and concentrator-5 Kit (Zymo research). After elution with 20 µL of nuclease-free water, the samples were incubated with 1 µL Exonuclease I (NEB) and 2 µL of 10× reaction buffer at 37 °C for 30 min, followed by enzyme inactivation at 80 °C for 20 min.

Double-strand (ds) cDNAs were generated by PCR amplification in 50 µL total reaction volume using the Advantage 2 PCR Enzyme System (Clontech). PCR reaction was performed using 20 µL cDNA from the previous step, 5 µL of 10× Advantage 2 PCR buffer, 1 µL of dNTPs 50×, 1 µL of 10 µM LA-oligo (Microsynt), 1 µL of Advantage 2 Polymerase and 22 µL of nuclease-free water following the program (95 °C—1 min, 11 cycles: 95 °C—15 s, 65 °C—30 s, 68 °C—6 min, 72 °C—10 min). Full-length double-stranded cDNA was purified with 30 µL of AMPure XP magnetic beads (Beckman Coulter), eluted in 12 µL of nuclease-free water and quantified using the dsDNA QuantiFluor Dye System (Promega).

The sequencing libraries were built by tagmentation using 50 ng of ds cDNA with the Illumina Nextera XT Kit (Illumina) following the manufacturer’s recommendations. The reaction was incubated for 5 min at 55 °C, immediately purified with DNA Clean and concentrator-5 Kit (Zymo research) and eluted with 21 µL of nuclease-free water. The tagmented library was PCR-amplified using 20 µL eluted cDNA, 2.5 µL of i7 Illumina Index, 2.5 µL of 5 µM P5-BRB primer (IDT) using the following program (72 °C—3 min, 98 °C—30 s, 13 cycles: 98 °C—10 s, 63 °C—30 s, 72 °C—5 min). The fragments ranging 300–800 base pairs (bp) were size-selected using SPRIselect (Beckman Coulter) (first round 0.65× beads, second 0.56×), with a final elution of 12 µL nuclease-free water. The resulting library was sequenced on an Illumina Hiseq 4000 sequencer as Paired-End 100 base reads following Illumina’s instructions. Image analysis and base calling were performed using RTA 2.7.7 and bcl2fastq 2.17.1.14. Adapter dimer reads were removed using DimerRemover (https://sourceforge.net/projects/dimerremover/).

### BRB-seq data processing

Pair-end reads with quality score higher than 10 were kept. The first read of the pair is 16 bases long. A first part of 6 bases corresponds to a unique sample-specific barcode and a second part of 10 bp is a unique molecular identifier (UMI). The second read of the pair, containing genomic data, was aligned to the human reference transcriptome from the UCSC website (release hg38) using BWA (*78*) (version 0.7.4.4) with the non-default parameter “−l 24”. Reads mapping to several positions in the genome were filtered out from the analysis. The pipeline is described in (*79*). After quality control and data pre-processing, a gene count matrix was generated by counting the number of unique UMIs associated with each gene for each sample.

### Immuno-histochemistry

Sections (5µm thick) of formalin-fixed, paraffin embedded tumor tissue samples were dewaxed, rehydrated through graded ethanol and subjected to heat-mediated antigen retrieval in citrate buffer (Antigen Unmasking Solution, Vector Laboratories). Slides were incubated for 10 min in hydrogen peroxide H2O2 to block endogenous peroxidases and then 30 min in saturation solution (Histostain, Invitrogen) to block nonspecific antibody binding. This was followed by overnight incubation with indicated primary antibodies at 4°C. After washing, sections were incubated with a suitable biotinylated secondary antibody (Histostain, Invitrogen) for 10 min. Antigen-antibody complexes were visualized by applying a streptavidin-biotin complex (Histostain, Invitrogen) for 10 min followed by NovaRED substrate (Vector Laboratories). Sections were counterstained with hematoxylin to visualize nucleus. Control sections were incubated with pool secondary antibodies without primary antibody. The antibody against the following target was used: CD34 (Abcam, ab81289).

### Immunofluorescence and histology image analysis

Formalin fixed paraffin embedded tissue blocks were sectioned at 5µm thickness. The tissue sections were dewaxed and rehydrated in xylene, gradual percentages of alcohols and finally in tap water. Protein epitopes of interest (CD8, particularly for CD8 positive lymphocytes and CD45 for all leukocytes) were retrieved in pH6 sodium citrate buffer using heat-induced epitope retrieval (HIER) method for 5 min, followed by quenching endogenous peroxidase activity and possible non-specific staining using 3% hydrogen peroxide (Sigma, H1009) and serum-free protein block (Agilent, X090930-2), respectively. Primary antibody - CD8 (Agilent, M710301-2, 1:800) and CD45 (Abcam, ab40763, 1:500) - was incubated for 1 hour at room temperature, followed by HRP (horseradish peroxidase) conjugated secondary antibody (Leica biosystems, DS9800) for 30 min. Fluorophore conjugated TSA (tyramide signal amplification) (Akoya bioscience, NEL744001KT, NEL741001KT) was used to visualize the primary antibody for 20 min. Then the sections were counterstained in Hoechst (Thermo fisher, H3570, 1:100) for 10 min and mounted in prolong gold anti-fade mounting buffer (Thermo fisher, P36930). Zeiss axio scan z1 was utilized to acquire digitized images using fluorescence channels such as Hoechst, FITC and Cy3. Images were then analyzed using HighPlex FL module in Halo AI (Indica Labs®version 3.6.4134) according to published protocol (*80*).

### Cell deconvolution algorithms

Prediction of tumor micro-environment (TME) cell type proportions from bulk gene expression data was performed using CIBERSORTx (version 1.0, 12/21/2019) and MuSiC (version 1.0.0) algorithms (*39, 40*). The CIBERSORTx cell fractions module was executed using the docker image after registration and access token received (https://cibersortx.stanford.edu/download.php). For both methods, reprocessed and relabeled single-cell RNA-seq data were used as reference (*34*). The 3 tumor predicted fractions (”Tumor cells”, “Cycling Tumor cells”, “Tumor cells CA9-high”) were summed into one “Tumor” fraction for clustering of samples based on estimated cell fractions.

### Pseudo-bulk RNA-seq mixtures

The generation of pseudo-bulk mixtures was performed from a single-cell RNA-seq data matrix made of several labeled cell types with gene expression in read counts. Step (1) consisted of sampling the single-cell RNA-seq data matrix. The number of cells of each cell type *i* used to construct the pseudo-bulk sample was defined using a random number *R* generated from Dirichlet’s law multiplied by the number of cells *S* of that type *i* present in the single-cell RNA-seq data matrix.

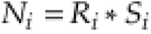

A matrix *M*, corresponding to the cells belonging to a pseudo-bulk mixture, is therefore made up of *N_i_* columns for each cell type *i*. Step (2) consisted in obtaining a pseudo-bulk sample by summing per gene the count values of reads across the cells of the different cell types. Steps (1) and (2) were repeated as many times as the number of pseudo-bulk samples to be generated.

The proportions of the different cell types composing each pseudo-bulk sample being known, the pseudo-bulk samples were used to evaluate the performance of the cell deconvolution algorithms. Comparisons between expected and predicted cell proportions were made using Spearman’s correlation coefficient, Root Mean Square Error (RMSE) coefficient and linear regression slope.

### Tumor purity analysis

A tumor purity score corresponds to the proportion of tumor cells in a sample. A tumor purity score was calculated for each sample from the gene expression values of its bulk transcriptome using the ESTIMATE method (Estimation of STromal and immune cells in MAlignant Tumor tissues using Expression data) (*42*). When available, the tumor purity score calculated from genomic profiling data of each tumor sample using the ABSOLUTE method (*41*) was also collected.

### Tumor-Immunity Differential score

The Tumor-Immunity Differential (TID) score reflects the difference between tumor and immunity activities in samples. It was built from four predicted cell fractions (Tumor, CD8.T, Plasma and Treg cells) and genomic features (Tumor Mutation Burden (TMB) score, *PDCD1* (PD-1) gene expression) frequently associated with patient response to ICB treatment. All features were standardized using Z-score transformation to make them comparable. The TID score was then defined as the difference between features considered unfavorable to the patient’s response to treatment, tumor cell fraction and TMB score (For the Braun et al. 2020 cohort of metastatic samples, 11 missing TMB scores were inferred by the median value), and those considered favorable, PD-1 expression and for regulatory T cell, CD8+ T cell and Plasma cell fractions (Eq. 1). The status of each feature was inferred from their independent association with patient clinical benefit.

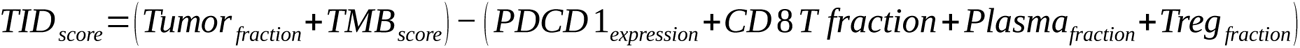

Equation 1: calculation of the Tumor-Immunity Differential score

Therefore, the higher the TID score, the less favorable the patient’s prognosis. High-TID and Low-TID sample categories were defined according to the median value of the TID score distribution.

### Clustering analysis

Unsupervised consensus clustering methods were implemented by the R/Bioconductor package ConsensusClusterPlus (version 1.62.0). Clustering of cell fractions and gene expression values were performed by k-means (parameters: reps = 1000, pItem = 0.8, pFeature = 1, clusterAlg = “km”, distance = “euclidean”). The best number of clusters was assessed by the delta area plot of consensus Cumulative Density Function (CDF). To construct relevant groups of ccRCC samples in terms of treatment response we retained only cell fractions predicted by the CIBERTSORTx algorithm associated with clinical benefit (two-sided Wilcoxon rank-sum test, corrected p-value < 0.05 or corrected p-value < 0.20, for, respectively, metastatic tumor and others if no one fraction was selected by the first threshold). The subtypes (C1-C3) of the 311 ccRCC samples with treatment response were found based on the consensus clustering of the selected cell fractions.

### Statistical tests

Statistical differences of cell fraction or gene expression values for comparisons were assessed using two-sided Wilcoxon rank-sum test with the Benjamini-Hochberg correction for multiple hypothesis testing. For sample comparisons across three or more groups, Kruskal-Wallis and two-sided chi-squared tests were used for numerical or categorical values, respectively.

Survival analyses were performed using the R package survival (version 3.4-0). The Kaplan-Meier curves of Progression-Free Survival (PFS) and Overall Survival (OS) were used to compare prognosis. The statistical comparison of the survival outcomes between subtypes was done using the log-rank test from the R package survminer (version 0.4.9).

The R package clusterProfiler (version 4.6.0) was used to perform over-representation enrichment analysis of the 6 differentially expressed genes (DEG) between C1-C3 clusters based on the Gene Ontology (GO) Biological Process database.

Single-sample Gene Set Enrichment Analysis (ssGSEA) of the hallmark (H) gene set from the MSigDB database (Human MSigDB v2023.1.Hs) was performed using the R/Bioconductor package GSVA (v. 1.42.0).

Single-sample Gene Set Enrichment Analysis (ssGSEA) was used to compute the MHCI and MHCII prognostic scores based on previously published gene signatures (*21*). Gene Set Variation Analysis (GSVA) was used to compute the MIAS and GEP prognostic scores based on gene signatures previously reported (*20, 22*). The JAVELIN score was calculated as the average of the standardized values of the 26 genes within the 26-gene JAVELIN Renal 101 Immuno signature (*1*). The IMPRES score was calculated from the method previously published (*19*). The two Tertiary Lymphoid Structure (TLS) scores were computed based on the mean of a 9-gene signature (CD79B, EIF1AY, PTGDS, RBP5, SKAP1, LAT, CETP, CD1D, CCR6) (*45*) or a 7-gene signature (CCL19, CCL21, CXCL13, CCR7, CXCR5, SELL, LAMP3) (*29*).

## List of Supplementary Materials

Present a list of the Supplementary Materials in the following format.

Materials and Methods

Fig S1 to S18 (on request)

Data file S1-S27 (on request)

## Acknowledgments

Most of the computations presented in this paper were performed using the GRICAD infrastructure (https://gricad.univ-grenoble-alpes.fr), which is supported by Grenoble research communities.

## Author contributions

Conceptualization: CB, HA, FJ, SS

Methodology: CB, FJ, SS, HA

Investigation: FJ, SS, PB, OF, CP, DH, IU

Visualization: FJ, SS

Funding acquisition: CB, HA, AB

Project administration: CB, HA

Supervision: CB, HA, DP

Writing – original draft: FJ, CB, SS, HA

Writing – review & editing: FJ, SS, CB, HA, DP, DH, OF, BE, ES, FC, JA, CP, FZ, MS, SNS, AL, JL, JD

## Data and materials availability

Processed bulk RNA-Seq and clinical data were collected for 5 different cohorts. A cohort of 331 primary and metastatic advanced clear cell Renal Cell Carcinoma (ccRCC) samples (EGAC00001001519; EGAC00001001520; EGAC00001001521) (*2*). A cohort of 11 primary or metastatic ccRCC samples (*49*) collected from (*22*). A cohort of 64 advanced melanoma samples from patients included in clinical trial CA209-038 (NCT01621490) treated with Nivolumab (GSE91061) (*51*). A cohort of 20 metastatic melanoma samples from patients treated by anti-PD-1 (GSE168204) (*52*). A cohort of 19 PBMC samples collected from blood of metastatic NSCLC patients (GSE111414) (*53*). Cell deconvolution results are provided in supplementary materials. Gene expression of ccRCC samples produced are available in EGA (). The code used to analyze cell deconvolution results and association with ICB data as well as the single-cell matrix refined used as reference in the cell deconvolution approach are available in Zenodo (ZenodoURL).

